# An inexpensive remotely-operated video recording system for continuous behavioral observations

**DOI:** 10.1101/596106

**Authors:** W.David Weber, Heidi S. Fisher

## Abstract

Video recording technology is an important tool for studies of animal behavior because it reduces observer effects and produces a record of experiments, interactions among subjects, and contextual information, however it remains cost prohibitive for many researchers. Here we present an inexpensive method for building a remotely-operated video recording system to continuously monitor behavioral or other biological experiments. Our system employs Raspberry Pi computers and cameras, open-source software, and allows for wireless networking, live-streaming, and the capacity to simultaneously record from several cameras in an array. To validate this system, we continuously monitored home-cage behavior of California mice (*Peromyscus californicus*) in a laboratory setting. We captured video in both low- and bright-light environments to record behaviors of this nocturnal species, and then quantified mating interactions of California mouse pairs by analyzing the videos with an open-source event logging software. This video recording platform offers users the flexibility to modify the specifications for a range of tasks and the scalability to make research more efficient and reliable to a larger population of scientists.

## Introduction

Behavior often represents an animal’s most immediate and dynamic response to internal or external cues [1]. Behavioral data are, therefore, invaluable indicators of change, yet the complexity and diversity of behaviors can be challenging to quantify. Traditional sampling methods from live observations (e.g., scan sampling, focal animal sampling, one-zero sampling [2–3]) frequently results in an estimate, rather than a precise measure of behaviors, or only provides data from a subset of subjects or interactions within a social group [4]. The behavioral record is therefore often incomplete with live observational data, and the presence of an observer can influence the behavior of study subjects, thus biasing the results [5–7]. Moreover, to fully understand an animal’s response, it is imperative to understand the context and the environment in which the behavior was expressed. Video recording enables researchers to review behaviors multiple times to yield a more complete dataset, reduces observational biases, and can capture contextual information.

Video recording has become a nearly ubiquitous tool for behaviorists and useful in a wide range of studies. For example, video can improve data collection in manipulative behavioral assays from studies of visual recognition in cichlids [8] and foraging in fruit flies [9], to parental care [10] and anxiety in rodents [11]. Moreover, passive monitoring approaches, such as home-cage cameras, can be used to record social behaviors, such as pair bonding in marmosets [12], or mating in whiteflies [13], guppies [14], and gerbils [15]. Outside of the lab, passive monitoring with nest box cameras have revealed unexpected behaviors including paternal care in tree swallows, even when extra-pair copulations were evident to the male [16], personality traits and social dominance in zebra finch [17], and long-term social interactions in “near natural” enclosures in house mice [4]. Continuous video recording systems provide a record of behavior and contextual information in a variety of studies from controlled laboratory assays to passive monitoring in the wild.

With few exceptions, continuous video recording improves the rigor and repeatability in behavioral studies, yet the technology remains cost prohibitive for researchers with limited funding or those requiring numerous cameras for complex experimental designs. Here we present an inexpensive method for building a remotely-operated video recording system using open-source software that allows for wireless networking, live-streaming, and the capacity to simultaneously record from several cameras to create an array. We then present validation of this system, in which we built an array of modules to continuously record mating behavior in California mice, *Peromyscus californicus*.

## Materials and Methods

We constructed a video recording array using Raspberry Pi (RPi; Caldecote, Cambridge, UK) modules and open-source software to continuously record multiple mating pairs of captive California mice (*Peromyscus californicus*) across several reproductive cycles. We then transferred the video files to Google Drive (Google LLC, Mountain View, CA, USA) for cloud-based storage, and later scored videos using the free open-source software, BORIS (Behavioral Observation Research Interactive Software [18]).

### Recording module assembly

Raspberry Pi computers are very small, inexpensive computers that offer a flexible platform to build a video recording system. Some base units can be purchased with cameras but storage and cables must be purchased separately. Table 1 describes the basic starting materials we used and ordering information. At the time of purchase, each recording module cost roughly $50.00 to build.

**Table 1.**
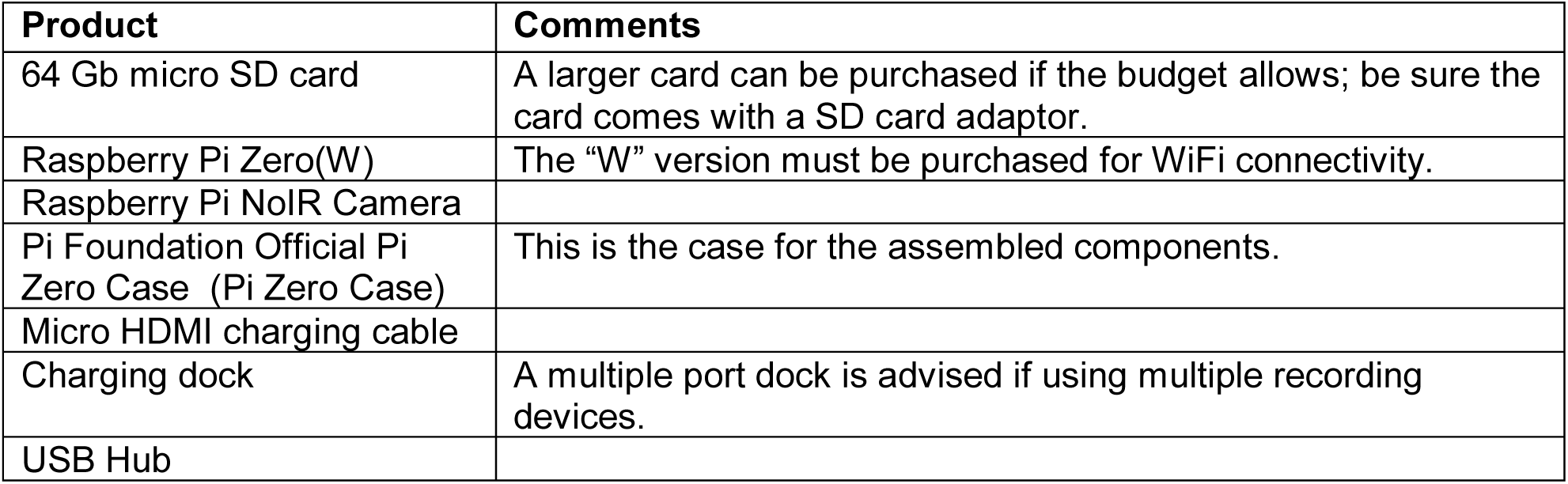
Basic Materials.

To build a recording module, we first loaded all application files, including an operating system (OS), onto a micro SD card. We used 64 Gb micro SD cards, as roughly 8 Gb must be allocated to the OS and our video files averaged 30 Gb in size. We formatted the card as MS-DOS(Fat); the “Raspbian” OS used in this application was designed to be loaded onto cards smaller than 32 Gb, therefore the 64 Gb card required reformatting. Once formatted, we loaded the “Raspbian” OS (https://www.raspberrypi.org/downloads/noobs/) onto the card and then installed it into the micro SD card port of a Raspberry Pi Zero(W) board (Figure 1A).

Next, we attached the Raspberry Pi NoIR camera to the R-Pi board (Figure 1B) and placed the board into the Pi Zero Case (Figure 1C). This step requires extreme care, as there are locking pins (see arrows in Figure 1D) that secure the board and camera to the case, thus it was necessary to bend the case to prevent damaging the R-Pi board. Once the case was closed, the recording module was complete (Figure 1D). We next attached the recording module to a monitor, mouse and keyboard with a USB hub, and then connected the module to a power source to activate the unit. The OS automatically initializes as soon as power is connected, the activation process runs for approximately 10 minutes and then the Raspian desktop appears.

**Figure 1.**
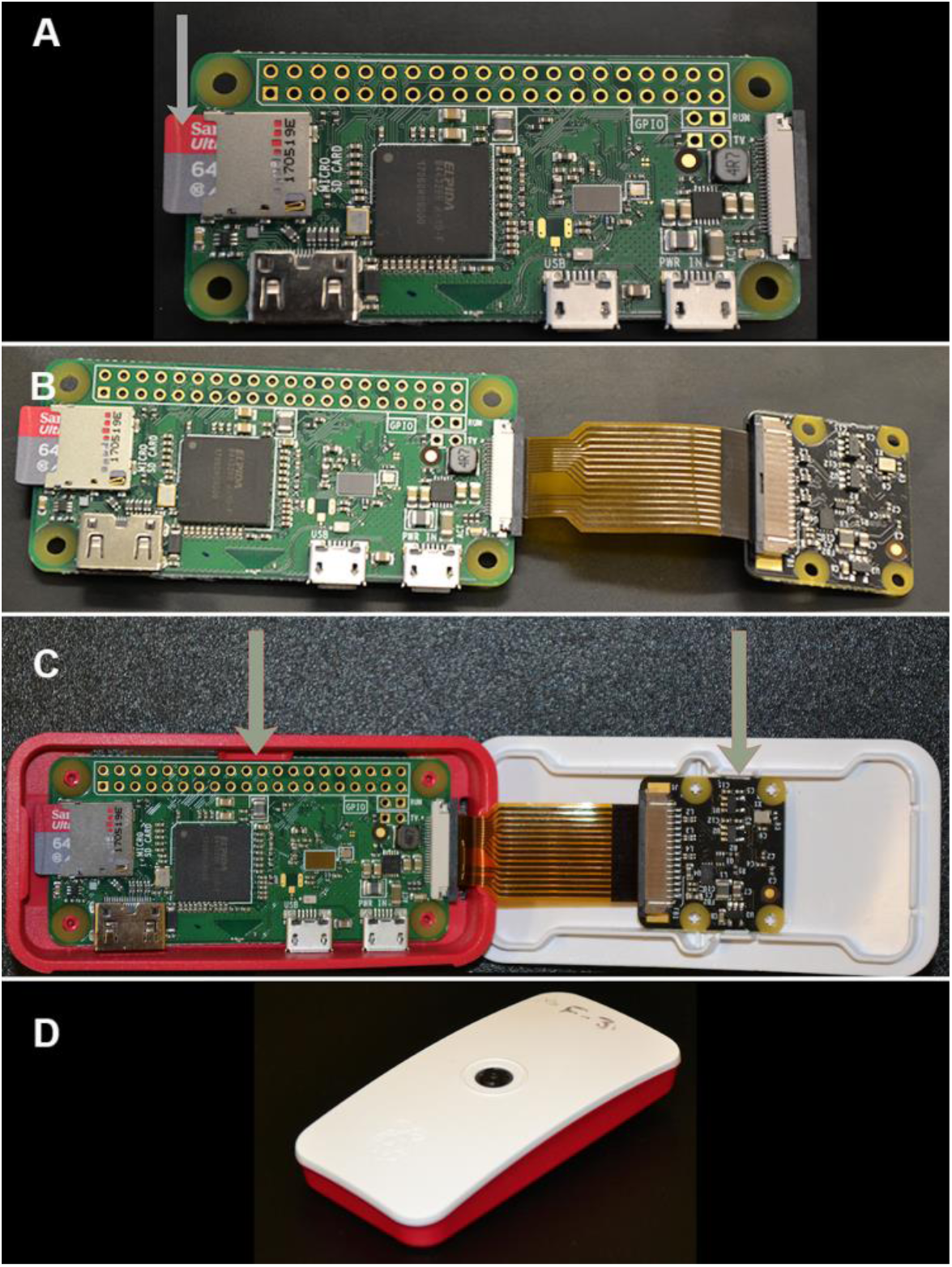
Assembly of a recording module. (A) MicroSD card installed into a Raspberry Pi Zero(W) board. Grey arrow indicates the location of the MicroSD card port. (B) Raspberry Pi board is attached to the camera assembly. (C) Pi Zero Case. Grey arrows indicate locking pins that the board and camera must fit underneath. (D) An assembled recording module.

### Recording Module Programming

On all modules, the default username is “pi” and password is “raspberry”, making them susceptible to unauthorized off-network access. Therefore, we next reset the default username and password in the “Raspberry Pi Configuration Menu”. Under the interface tab of that same menu, we then enabled the “Camera”, selected “SSH” (allows for remote access) and “VNC” (allows for remote interfacing) options, and then rebooted the module. After reboot, we established an internet connection by hovering the cursor over the WiFi icon, selected the appropriate network, and logged in. By hovering the cursor over the WiFi icon after establishing a connection, the IP address is revealed; with this IP address we could remotely log into the module using RealVNC Viewer (https://realvnc.com/en/), an open-source software.

### Behavioral Observations

To assess the routine use of the video module to record social interactions, we observed mating behavior in California mice. We obtained four male and four female mice from the Peromyscus Genetic Stock Center at the University of South Carolina (https://www.pgsc.cas.sc.edu/), and housed them at the University of Maryland, in accordance with the Institutional Animal Care and Use Committee policies (protocol number: R-Jul-18-38), and in standard transparent laboratory rodent cages lined with cedar shavings. We provided food and water *ad libitum* and maintained all mice at 22°C in a 16:8 light-dark cycle, simulating this species’ breeding season [19].

To record the mating behavior, we placed a male and a female into a cage on a camera rack (Figure 2). Camera stands can be easily purchased online or 3D printed, but we fashioned a rack from metal shelving on which we attached two blocks cut from recycled styrofoam at either end to hold the two recording modules per cage, in the middle of which we placed a cage (Figure 2). We recorded from four pairs, each with their own pair of recording modules. We recorded behaviors during the light period with one module and used a second module to record behavior during the dark period while the room was illuminated with red lights and infrared LED lights. We found that using two modules, each set to parameters optimized for each light environment, was the most effective way for us to obtain high quality video across the entire light-dark cycle, however a single camera can be programed to adjust parameters at different times through the day. In our setup, the size of the files produced from using a single module exceeded the capacity of the micro-SD card, and we found that adding as second module was more cost effective that using a high-capacity micro-SD card.

**Figure 2.**
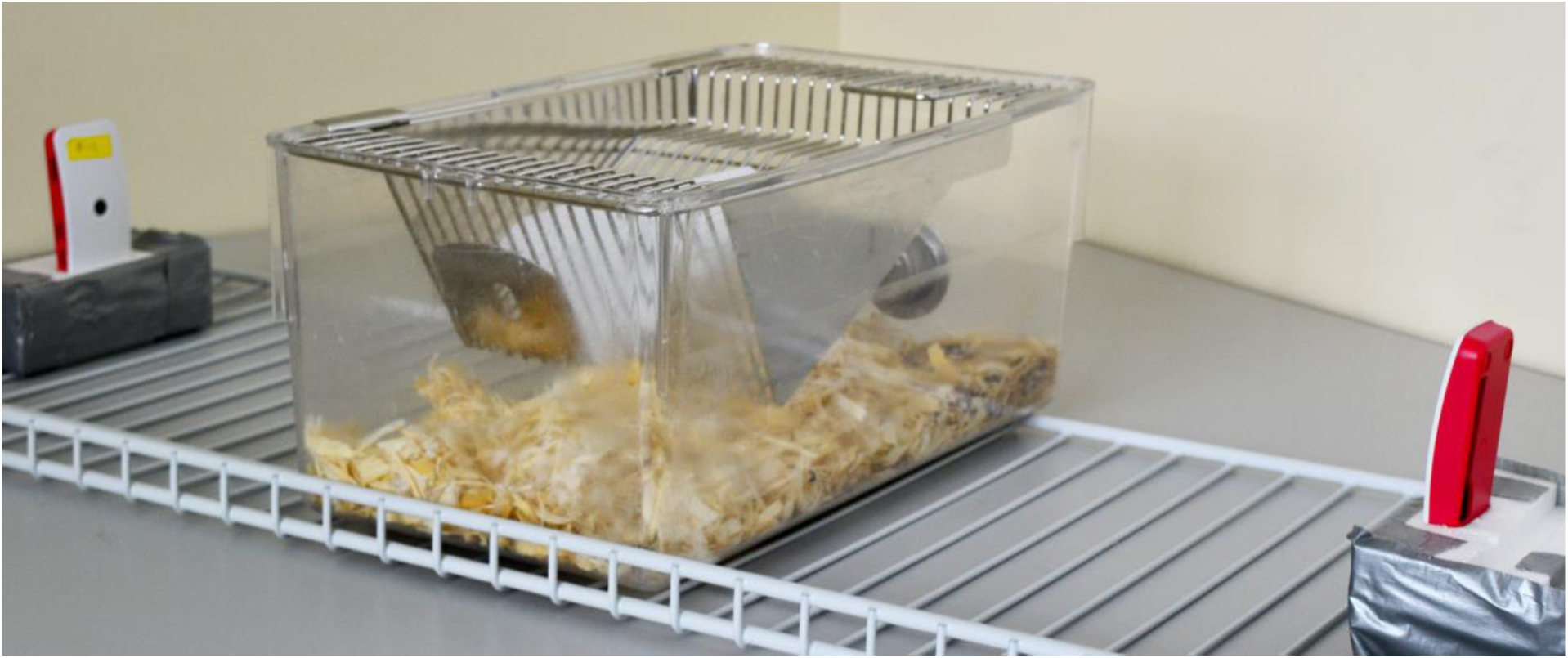
Assembled camera rack with animal cage.

Next, we remotely logged into the modules using a Python 3 shell (selected from the programming menu on the Raspbian desktop) and executed the camera script (Table S-1; all coding scripts described in this article can also be accessed at https://github.com/wdavidweber/Pi-Recording-Module). To send video files directly to a Google Drive folder, we used an additional script because the software is not included in the Raspian OS. Using the terminal console in Raspbian, we uploaded the “rclone” (Nick Craig-Wood ©2012) open-source software (Table S-2) and regularly monitored the modules and Google Drive folders. Free storage on Google Drive is limited to 15Gb, unless an unlimited account is available to you, therefore, files may need to be downloaded and transferred prior to filling the storage. To score the behaviors in BORIS, we converted the .h264 format files (the default Raspbian filetype) into “.mp4” format using the “ffmpeg” package, available in the Raspbian OS (the code for this is the last line of script in Table S-1). We performed the file conversion using a High-Performance Computing Cluster at the University of Maryland because our video files were too large and numerous to be converted on the modules. In most situations, this can be done on the module itself, or the conversion can be performed on any workstation. When necessary, we live-streamed footage with a YouTube account (https:www.youtube.com/) by loading “Docker” onto the module (Table S-3). With this method, video recording and live-streaming cannot be accomplished simultaneously.

To quantify mating behavior, we scored videos using the open-source software, BORIS, and developed an ethogram based on the work of Dewsbury [20]. We recorded the frequency of mounts, intromissions, and ejaculations, as well as intromission latency (the time between the start of a reproductive bout and the first intromission), ejaculation latency (the time between the first intromission and the subsequent ejaculation), post-ejaculatory interval (the time between an ejaculation and the beginning of the next copulatory bout), reproductive event (the total duration of all mating events during a female’s single reproductive period, beginning with the first mounting event and concluding with the last ejaculation), copulatory bout (the duration between the first mounting event and the first ejaculation).

## Results

We continuously recorded home-cage behavior for three weeks from four pairs of California mice. In total, we collected 2016 hours of video, which resulted in roughly 2 TB of data. Recording one hour of footage during the light period (Figure 3A-B) under white lights, at a resolution of 1280 × 768, a framerate of 15 frames per second, and a brightness value of 70, resulted in recordings that were 440 Mb to 500 Mb in size. Recording one hour of footage during the dark period (under room red lights and LED infrared lights mounted above the cage; Figure 3C-D) at a resolution of 1280 × 768, a framerate of 15 frames per second, and a brightness value of 80, resulted in recordings of 1.5 Gb to 2 Gb in size. This difference in brightness variables was the result of room illumination, there is more illumination in the room under white light than while under red light. By reducing the parameters for resolution, framerate, or brightness small files are generated, but the footage is less resolved.

**Figure 3.**
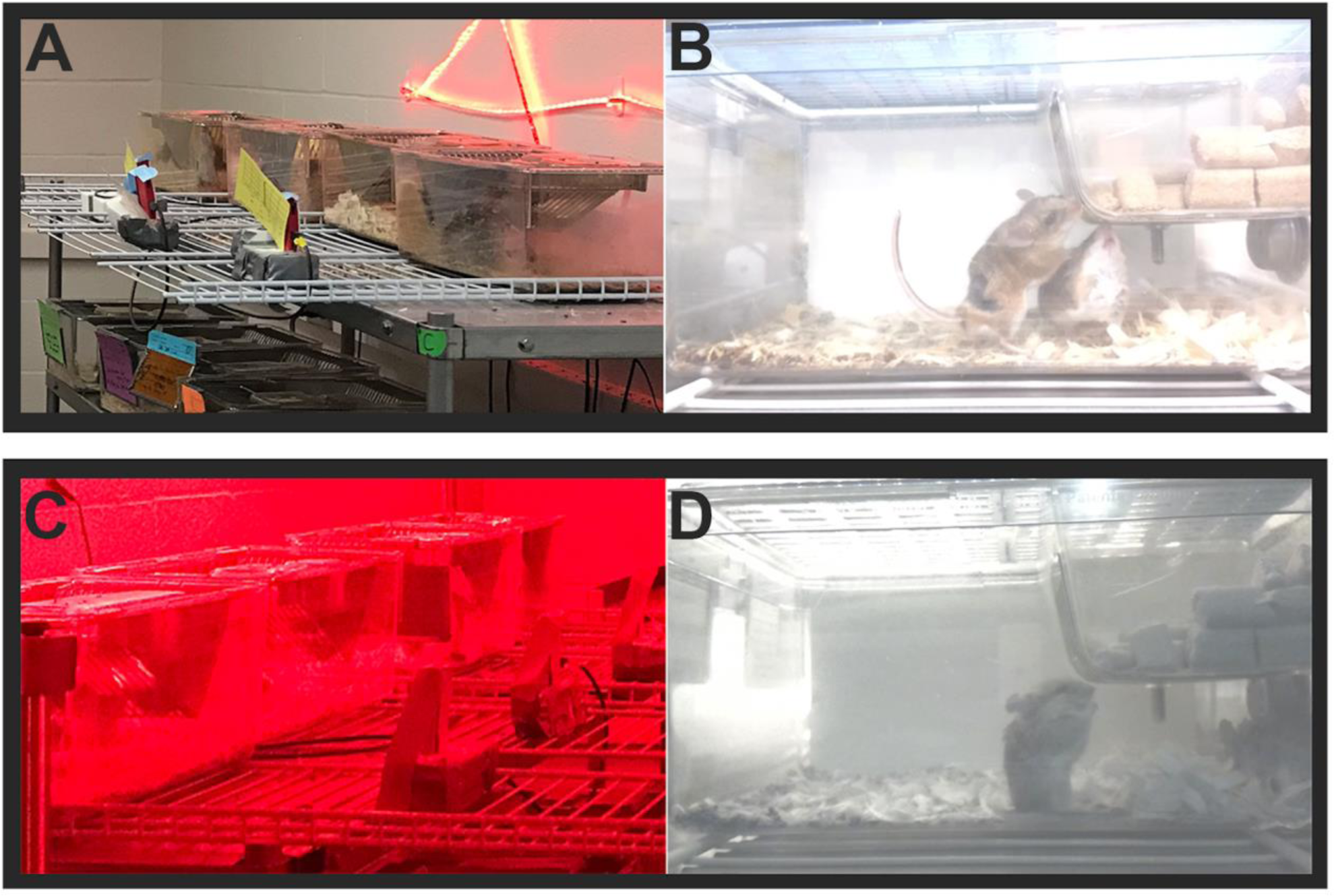
Recording module array and output. (A) The array during the light period with room illuminated by white lights, note the infrared LED lights over the cages (not needed for bright light conditions, shown here as an example). (B) Still of a video obtained during the light period, one animal as a spot shaven on his back for identification purposes. (C) The same array in (A) but the image is taken from the opposite side while during the dark period under room red lights and infared LED lights mounted above cages (not shown here). (D) Still image from a video obtained during the dark period.

We scored mating behavior from the one pair that successful produced a litter after the mating trial. The active mating period spanned a total of 5 hours, 24 minutes, and 3 seconds of footage, all of which occurred during the dark period of the photocycle and included two copulatory bouts. The first mating bout spanned 4 minutes, 15 seconds, and included 8 mounts, 3 intromissions and a single ejaculation; the second bout spanned 2 minutes, 45 seconds and included 6 mounts, 5 intromissions and a single ejaculation. We found that the mean intromission latency was 1 minute, 49 seconds and the mean ejaculation latency was 1 minute 42 seconds. The two copulatory bouts were separated by a 36 minutes, 39 second refractory period.

## Discussion

Here we describe a remotely-operated video recording system to monitor animal behavior using wireless networking. This tool permits user-determined specifications, including live-streaming and multiplexing, and has the capacity to simultaneously record and upload data files for storage. Moreover, this system allows for continuous behavioral monitoring on a flexible timescale. The cost of the recording module that we describe is less or comparable to some pre-assembled commercial surveillance cameras, which like the RPi recording module, often use SD cards to store footage and may provide cloud storage for a fee with restricted allocations. However the system described here offers a wider range of low-cost modifications to suit a variety of research environments by incorporating open-source software add-ons.

For validation purposes, we used the video capture system we report here to record, and subsequently quantify, mating behavior in the California mouse. Once we positioned the cameras in place in our animal facility, we controlled the modules from our laboratory located in another building. This design eliminated possible observer effects on the animals’ behavior, provided contextual information on the animals’ environment, and allowed for off-hours monitoring of the animal housing space. Furthermore, with the addition of infrared LED lights, we were able to record mating behavior of this nocturnal species without interruption or disturbance.

In the research setting, our video recording system can be applied to tasks beyond animal behavior experiments. For example, this system could be used to remotely monitor chemical reactions or other assays in which timing is uncertain, or if the modules are housed within an incubator, to monitor bacterial or embryo culture. The flexible platform, scalability and low cost of our “do it yourself” video recording system is a powerful lab tool to make research more efficient and reliable to a larger population of scientists.

## Acknowledgements

We thank Andrés Bendesky, whose use of Raspberry Pi cameras to record parental care behavior in *Peromyscus* inspired us to build and report on the system presented here. We also thank Irene Lui for her computer programming guidance, Erica Glasper for the experimental animals, and R. Zaak Walton, whose thoughtful discussion and advice was essential to this project.

## Supplemental Material

**Table S-1.** Camera operation and recording script

| Line | Python Script |
| --- | --- |
| 1 | `from picamera import PiCamera` |
| 2 | `from time import sleep` |
| 3 | `import os` |
| 4 |  |
| 5 | `camera = PiCamera()` |
| 6 |  |
| 7 | `camera.resolution = (1280, 768)` |
| 8 | `camera.framerate = 15` |
| 9 | `camera.brightness = 70` |
| 10 | `camera.start_preview()` |
| 11 | `camera.start_recording('/home/pi/Videos/Recording.h264')` |
| 12 | `sleep(61200)` |
| 13 |  |
| 14 | `os.system('ffmpeg -vcodec h264 -i Recording.h264 -vcodec copy -acodec copy Recording.mp4')` |
| 15 | `os.system('rclone sync /home/pi/Videos/Recording.mp4 picam:AA')` |
| 16 | `rm /home/pi/Videos/Recording.h264` |

Explanation: (1) This line activates the software for the camera. (2) This line tells the unit to use the sleep software for timing. (3) This command allows the OS to operate in the python shell. (4) Skip this line. (5) This line instructs the camera to execute the following script. (6) Skip this line. (7) This sets the resolution for the recording. (8) This sets the framerate for the recording. (9) This sets the brightness for the recording. (10) This command begins camera operations at the script execution. (11) This command begins recording at the execution, and places files where directed [e.g., ("")]. (12) This command sets the recording time (e.g., 61200 seconds). (13) Skip this line. (14) This line instructs the R-Pi to convert the recording into .mp4 format. (15) This command uploads the recording onto the specified Google Drive folder. (16) This removes the .h264 file that was used to make the .mp4 video.

**Table S-2.**
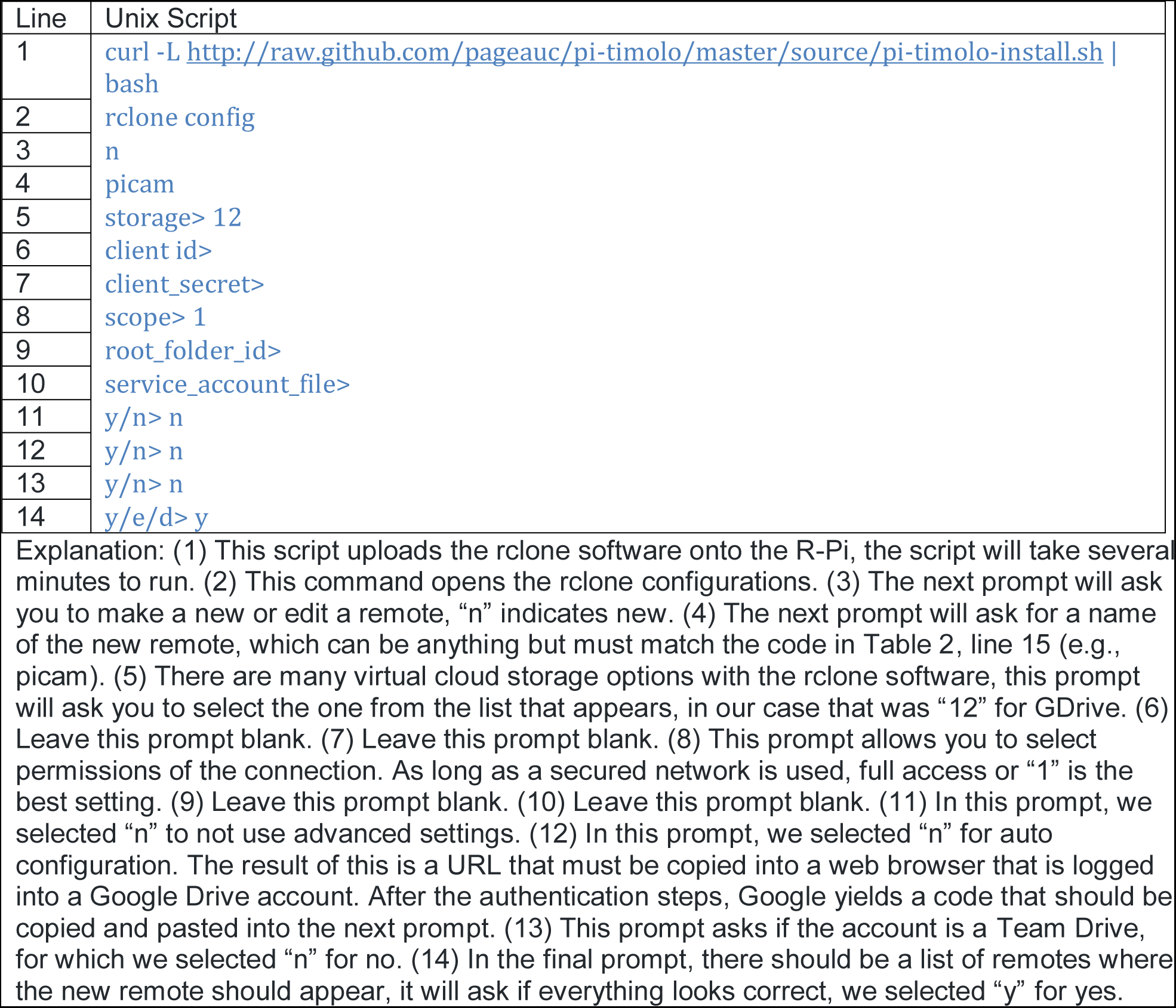
Rclone installation and GDrive connection script

**Table S-3.**
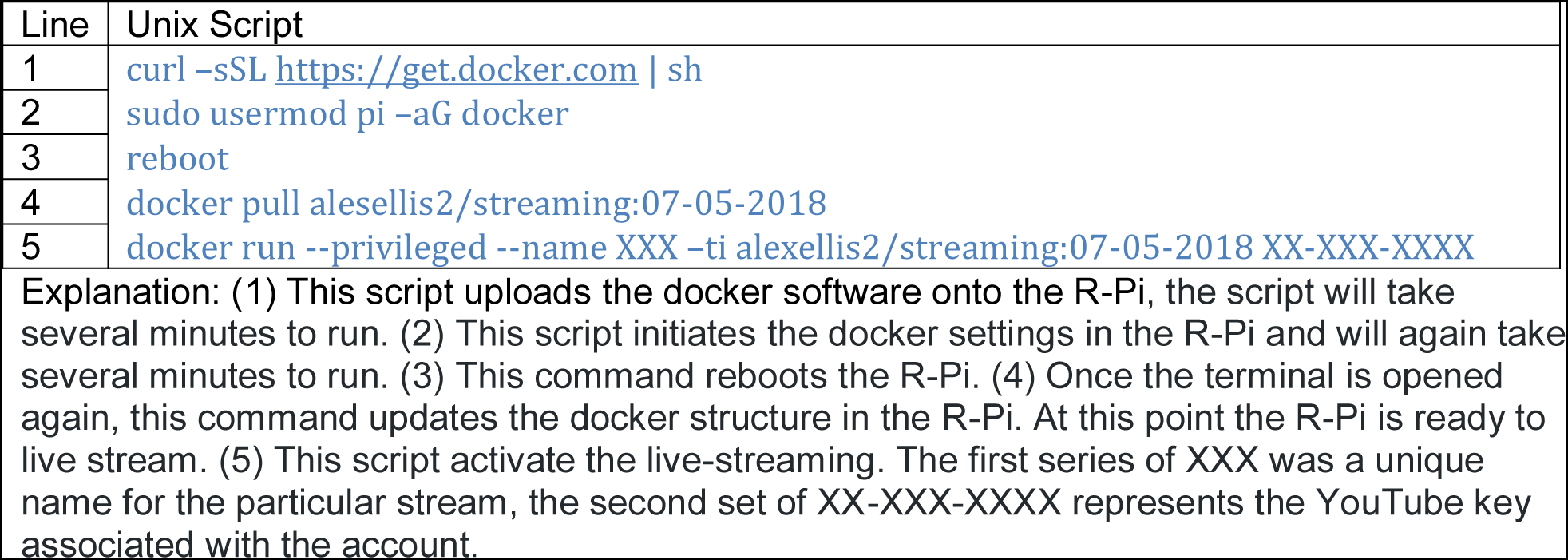
Docker script for live-streaming

## References

1. Bretman A, Gage M, Chapman T. Quick-change artists: male plastic behavioural responses to rivals. Trends Ecol Evol. 2011;26(9):467–73.

2. Altmann J. Observational Study of Behavior : Sampling Methods. Behaviour [Internet]. 1974;49(3/4):227–67.

3. Martin P, Bateson P. Measuring behaviour: An introductory guide. 2nd, editor. Cambridge, EN, UK: Cambridge University Press; 1995.

4. Weissbrod A, Shapiro A, Vasserman G, Edry L, Dayan M, Yitzhaky A, et al. Automated long-term tracking and social behavioural phenotyping of animal colonies within a semi-natural environment. Nat Commun. 2013;4(May):1–10.

5. Wilson S. The use of ethnographic techniques in educational eesearch. Rev Educ Res. 1977;47(2):245–65.

6. Monahan T, Fisher J. Benefits of “observer effects”: Lessons from the field. Qual Res. 2010;10(3):357–76.

7. Herbert R, James F. The Hawthorne experiments: first statistical interpretation. Am Sociol Rev. 1978;43(5):623–43.

8. Escobar-Camacho D, Marshall J, Carleton K. Behavioral color vision in a cichlid fish: *Metriaclima benetos*. J Exp Biol [Internet]. 2017;220(16):2887–99.

9. Jang EB, Light DM. Behavioral responses of female oriental fruit flies to the odor of papayas at three ripeness stages in a laboratory flight tunnel (*Diptera: Tephritidae*). J Insect Behav. 1991;4(6):751–62.

10. Bendesky A, Kwon YM, Lassance JM, Lewarch CL, Yao S, Peterson BK, et al. The genetic basis of parental care evolution in monogamous mice. Nature

11. Leuner B, Glasper ER, Gould E. Sexual experience promotes adult neurogenesis in the hippocampus despite an initial elevation in stress hormones. PLoS One. 2010;5(7).

12. Gerber P, Schnell C, Gusti A. Behavioral and cardiophysiological responses of Common Marmosets (*Callithrixjacchus*) to social and environmental changes. Primates. 2002;43(July):201–16.

13. Ruan YM, Luan JB, Zang LS, Liu SS. Observing and recording copulation events of whiteflies on plants using a video camera. Entomol Exp Appl. 2007;124(2):229–33.

14. Pilastro A, Mandelli M, Gasparini C, Dadda M, Bisazza A. Copulation duration, insemination efficiency and male attractiveness in guppies. Anim Behav. 2007;74(2):321–8.

15. Prates EJ, Guerra RF. Parental care and sexual interactions in Mongolian gerbils (*Meriones unguiculatus*) during the postpartum estrus. Behav Processes. 2005;70(2):104–12.

16. Whittingham L, Dunn P, Robertson R. Confidence of paternity and male parental care: an experimental study in tree swallows. Anim Behav. 1993;139–47.

17. Morgan D, Auclair Y, Cezilly F. Personality predicts social dominance in female zebre finches, *Taeiopygia guttata*, in a feeding context. Anim Behav. 2011;81(1):219–24.

18. Friard O, Gamba M. BORIS: a free, versatile open-source event-logging software for video/audio coding and observations. Methods Ecol Evol. 2016;7(11):1325–30.

19. Kirkland GL, Layne JN. Advances in the study of Peromyscus (Rodentia). Lubbock, TX, USA: Texas Tech University Press; 1989.

20. Dewsbury D. Copulatory behavior of White-Footed mice (*Peromyscus leucopus*). J Mammal. 1975;56(2):420–8.

